# Transient deSUMOylation of IRF2BP proteins controls early transcription in EGFR signaling

**DOI:** 10.1101/819730

**Authors:** Sina V. Barysch, Nicolas Stankovic-Valentin, Samir Karaca, Judith Doppel, Thiziri Nait Achour, Carsten Sticht, Henning Urlaub, Frauke Melchior

## Abstract

Molecular switches are essential modules in signaling networks and transcriptional reprogramming. Here, we describe a role for <u>s</u>mall <u>u</u>biquitin-related-<u>m</u>odifier SUMO as a molecular switch in epidermal growth factor receptor (EGFR) signaling. Using quantitative mass spectrometry, we compared the endogenous SUMO-proteomes of Hela cells before and after EGF-stimulation. Thereby, we identified a small group of transcriptional co-regulators including IRF2BP1, IRF2BP2 and IRF2BPL as novel players in EGFR signaling. Comparison of cells expressing wildtype or SUMOylation deficient IRF2BP1 indicated that transient deSUMOylation of IRF2BP1 is important for appropriate expression of immediate early genes including *Dual specificity phosphatase 1* (DUSP1, MKP-1), an important feedback regulator of EGFR signaling. We find that IRF2BP1 is a SUMO-dependent repressor, whose transient deSUMOylation on the DUSP1 promotor allows - and whose timely reSUMOylation restricts - DUSP1 expression. Our work thus provides a paradigm how comparative SUMO proteome analyses serve to reveal novel regulators in signal transduction and transcription.

## Introduction

Small ubiquitin related modifier (SUMO) is an essential protein modification that regulates hundreds of proteins and numerous processes including signal transduction and transcription processes (Gareau & Lima, 2010; Flotho & Melchior, 2013; Seeler & Dejean, 2017; Zhao, 2018). The last decade has seen a dramatic improvement of SUMO proteomics (as reviewed in (Hendriks & Vertegaal, 2016)), which culminated in several thousand SUMO-modified proteins and the startling number of 14,869 different SUMO2/3 acceptor sites in human cells during stress (Hendriks *et al*, 2018). Together with studies that revealed simultaneous SUMOylation of multiple subunits in protein complexes, e.g. upon DNA damage (Psakhye & Jentsch, 2012), and with the growing evidence that SUMO can contribute via low-affinity/high avidity interactions to phase separation (reviewed (Zhao, 2018)), this may lead to the impression that SUMO largely functions as a "spray" with little specificity. However, there are numerous examples where reversible SUMOylation of a single protein on a specific lysine residue determines protein function in highly controlled manner. A famous example is yeast PCNA, which interacts with the helicase Srs2 specifically upon S-phase specific SUMOylation (Pfander *et al*, 2005). But how can we move from lists of thousands of SUMO targets to those that are most relevant? We speculated that targets whose SUMOylation changes in response to a physiological stimulus may be of particular importance.

Comparative quantitative phospho-proteomic screens have been used successfully as discovery tools to identify important players, cancer drug targets or signaling branches, e.g. by comparing different types of cancers, wildtype versus mutant EGFR cells, or stimulation with platelet-derived growth factor PDGF versus EGF (reviewed in (Kolch & Pitt, 2010)). We thus reasoned that comparative analysis of SUMO, acting as a similar molecular switch, might be another tool to identify new key factors in signaling and signal-dependent transcription. Recently, we developed a method that allows identification and quantitative comparison of endogenously SUMOylated proteins (Becker *et al*, 2013; Barysch *et al*, 2014). This method is very well suited to identify individual proteins that may alter their SUMOylation status in response to a physiological stimulus.

For a "proof of principle" study, we turned to epidermal growth factor receptor (EGFR) signaling in Hela cells. EGFR signaling is one of the most prominent, best-characterized and essential signaling networks in metazoans. It is involved in cellular growth and development, as well as in cancer progression. Key to its many roles is the transcriptional reprogramming of cells. After stimulation with EGF, a highly complex signaling network is activated, several actors work in parallel and compensatory mechanisms as well as feedback-loops are installed (Kolch & Pitt, 2010; Citri & Yarden, 2006). In the so-called early loops, phosphorylation and ubiquitylation play essential roles to control ligand-induced receptor endocytosis and cytosolic signaling events. The late loops involve transcriptional regulation, which acts in three temporal phases, the immediate early gene (IEG), the delayed early gene (DEG) and the secondary response gene (SRG) transcription waves (Avraham & Yarden, 2011).

Large quantitative studies have been performed to investigate EGF-induced changes in the phospho-proteomes (Kratchmarova *et al*, 2005; Olsen *et al*, 2006; Oyama *et al*, 2009) and the ubiquitin-proteome (Argenzio *et al*, 2011). Those studies led to the identification of important regulatory events, mainly in the early loop of the EGF response. Here we investigated changes in the endogenous SUMO1 and SUMO2/3 proteome in EGFR signaling using quantitative mass spectrometry 10 min after- or without EGF treatment. As detailed below, we could indeed identify a SUMO-dependent molecular switch in EGF receptor signaling that involves the IRF2BP family of transcription co-regulators. IRF2BP proteins gain increasing interest, particularly in the context of inflammation, but they have not yet been linked to EGF receptor signaling. Our findings reveal that this protein family plays an important role in immediate early gene expression and feedback regulation, and demonstrate how transient deSUMOylation of a repressor can serve to control temporal gene expression.

## Results

### EGF stimulation induces transient deSUMOylation of several transcriptional regulators

To address the question whether EGFR signaling induces changes in the SUMO proteome and whether we are able to identify potentially new key factors in EGFR signaling, we combined our previously published method to enrich endogenously SUMOylated proteins (SUMO-IP, (Becker *et al*, 2013; Barysch *et al*, 2014)) with SILAC-based quantitative mass-spectrometry. For this, labeled Hela cells were serum-starved and treated with 0 or 100 ng/ml EGF for 10 min, respectively. This early time point was chosen to possibly detect both early events, e.g., at the plasma membrane, as well as early downstream events in transcription. Combined cell lysates were subjected to SUMO1- and SUMO2/3-IPs, followed by quantitative mass spectrometry in three independent experiments with charge swapping (Fig 1A). The vast majority of identified SUMO candidates (1228 for SUMO1 and 855 for SUMO2/3) were equally abundant in EGF treated and untreated samples. While no protein seemed to quantitatively lose or gain SUMO after 10 min of EGF-stimulation, 11 proteins could be identified whose abundance differed significantly (p<0.001) between both samples (Fig 1B and 1C, left panel, Dataset EV1). Intriguingly, five of these were transcriptional repressor proteins, TRIM24/TIF1α (Le Douarin *et al*, 1998) and TRIM33/TIF1γ (Venturini *et al*, 1999), as well as IRF2BP1, IRF2BP2 and IRF2BPL (Childs & Goodbourn, 2003). To validate this finding, we repeated the SUMO IPs and tested candidates by immunoblotting (Fig 1C, right panel). Indeed, as indicated by the decreased mobility of proteins in the IP compared to the input, IRF2BP1, IRF2BP2 and TRIM24 were SUMOylated and lost SUMO upon EGF treatment. RanGAP1 and TRIM28, two known SUMO targets whose abundance did not alter in the SUMO proteomic analysis, served as controls (Fig 1C). We next analyzed the kinetics of deSUMOylation by performing an EGF time course experiment. Surprisingly, deSUMOylation of TRIM24, IRF2BP1 and IRF2BP2 is very transient with a minimum after 15 min EGF-stimulation and full recovery after 60 min (Fig 1D).

**Figure 1:**
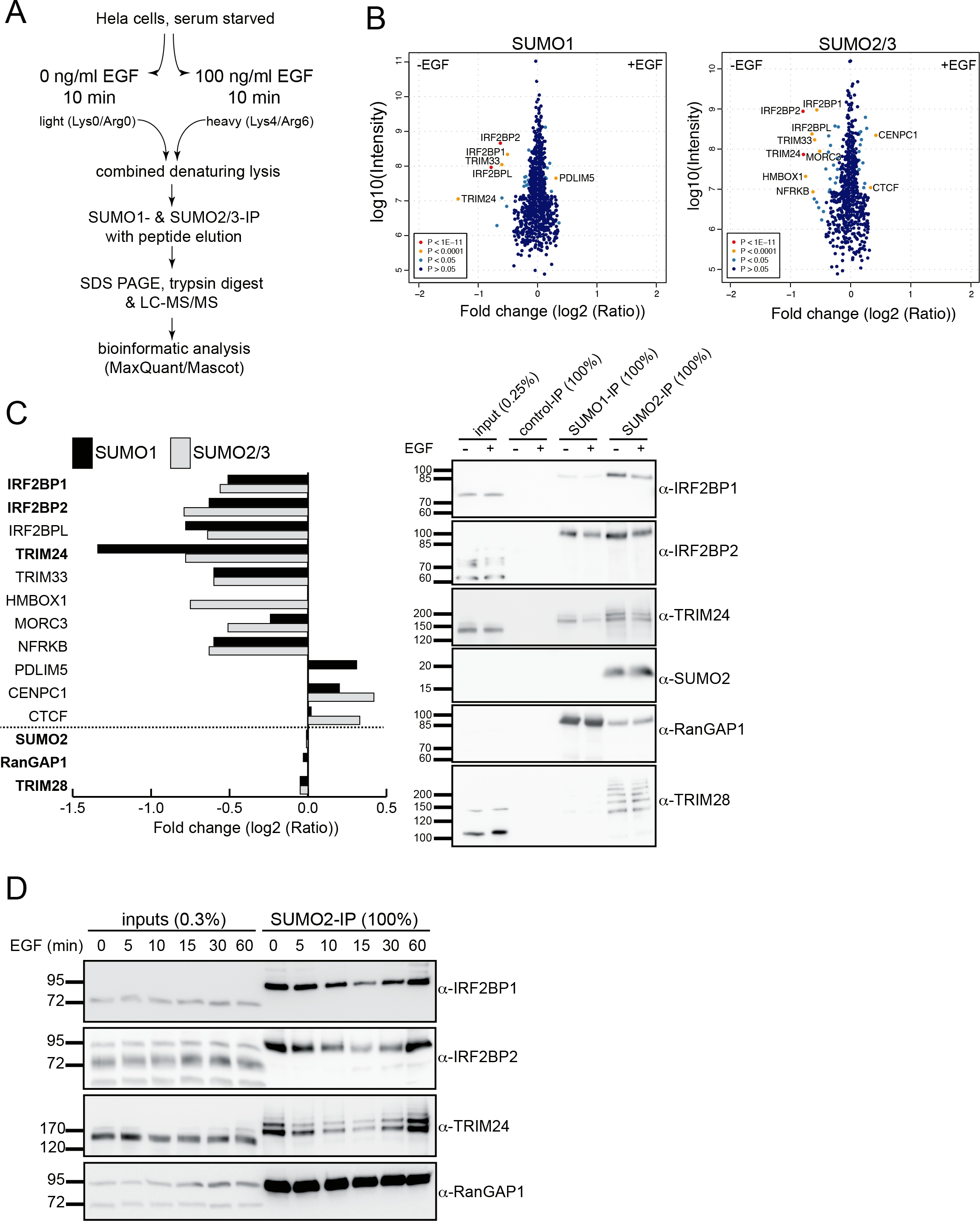
EGF induces rapid and transient deSUMOylation of transcriptional regulators in Hela cells. (A) Schematic representation of a quantitative proteome analysis that compares the SUMO proteome of serum-starved Hela cells treated for 10 min with or without 100 nM EGF (SILAC labeling and endogenous SUMO1- and SUMO2/3-IPs). (B) Scatterplots represent SILAC/SUMO-IP quantification after EGF treatment from three biological replicates. Each dot represents a protein that is either present in the same ratio between untreated and EGF treated cells (dark blue dots) or is significantly more present in one of the samples (red and yellow dots). (C) Left panel: Bar graph depicting mass spectrometry results of 11 proteins with altered SUMOylation (significant hits with P<0.0001), as well as the three non-changing proteins SUMO2, RanGAP1 and TRIM28. Right panel: Proteins highlighted in bold in the left panel were validated by SUMO-IP/immunoblotting from serum starved Hela cells without or with 10 min EGF stimulation. (D) Time course experiment: serum starved Hela cells were treated with 100 nM EGF, samples were harvested at indicated times and subjected to SUMO2 IP followed by immunoblotting with the indicated antibodies. IRF2BP1, IRF2BP2 and TRIM24 are rapidly but transiently deSUMOylated upon EGF treatment.

None of our five strongest hits has previously been described to be directly involved in EGF receptor signaling, even though TRIM24 can cross-talk with PI3K/AKT signaling (Zhang *et al*, 2015; Lv *et al*, 2017) and TRIM33, as well as IRF2BP1 and IRF2BP2 are known players in TGF-β signaling (Faresse *et al*, 2008; Quéré *et al*, 2014; Xi *et al*, 2011; Yuki *et al*, 2019; Kaiser Manjur *et al*, 2019). Furthermore, while TRIM24 and TRIM33 have been shown to be SUMOylated (Seeler *et al*, 2001; Fattet *et al*, 2013), SUMOylation of IRF2BP1, IRF2BP2 and IRF2BPL had not been investigated yet (see below).

### IRF2BP proteins are SUMOylated at their conserved C-termini

IRF2BP proteins are transcriptional co-regulators that can homo- and hetero-oligomerize via a conserved N-terminal zinc finger and can interact with diverse transcription factors via a C-terminal RING domain (Childs & Goodbourn, 2003; Yeung *et al*, 2011). They have initially been identified as putative transcriptional repressors in a yeast two hybrid screen as interaction partners of IRF2 (Childs & Goodbourn, 2003). In the following years, several groups have described individual target genes that they transcriptionally regulate, such as the TGF-β-Smad target genes ADAM12 and p21Cip1 (Faresse *et al*, 2008), as well as VEGFA (Teng *et al*, 2010) and FASTKD2 (Yeung *et al*, 2011), to name some examples. More recently, the IRF2BP proteins have emerged as important factors in the immune system. They can suppress inflammation in macrophages and microglia (Chen *et al*, 2015; Cruz *et al*, 2017; Hari *et al*, 2017) and have inhibitory effects on the expression of the programmed death ligand 1 (PD-L1) (Dorand *et al*, 2016; Soliman *et al*, 2014; Wu *et al*, 2019). Furthermore, IRF2BP2 was found to regulate the Hippo pathway in liver cancer and to act as a tumor suppressor (Feng *et al*, 2019). A recently published ChIP-Seq study of IRF2BP2 in mouse MEL cells revealed more than 11.000 binding sites in the mouse genome (Stadhouders *et al*, 2015), 40% of which are found at proximal promoter regions (< 5 kb from TSS).

To investigate functional consequences of EGF-dependent deSUMOylation of IRF2BP proteins, we sought to identify their SUMO acceptor sites. SUMO modifies its target proteins with the help of SUMO-specific E1, E2 and E3 enzymes on lysine residues that are typically embedded in SUMO consensus sites (ΨKxE/D, where Ψ is a bulky hydrophobic residue), as reviewed in (Flotho & Melchior, 2013; Gareau & Lima, 2010). When we inspected IRF2BP1 and IRF2BP2 for putative SUMOylation sites, we found two motifs that were conserved within the family and between species, a minimal KxE motif (FKKD/E) in the otherwise poorly conserved middle region and a classic SUMO consensus site (VKKE) close to the C-terminus (Fig 2A). To test whether one of those predicted IRF2BP SUMOylation sites is indeed the functional one, we transiently transfected HA-tagged wildtype and KR-mutant variants of both proteins into Hela cells and performed endogenous SUMO-IPs. While SUMOylated IRF2BP wildtype and K247R or K326R mutants can be immunoprecipitated, no signal was detected for the C-terminal K579 or K566R mutants. Thus, the lysine within the predicted C-terminal consensus SUMO site seems to be the dominant one endogenously used in mammalian cells (Fig 2B). This SUMOylation site is located at the very C-terminus of IRF2BP proteins, within a stretch of 30 highly conserved amino acids that follow directly after the conserved RING domain (Fig 2C). Of note, this SUMOylation site was not identified in recent proteomic screens from human cells (Hendriks *et al*, 2014; Xiao *et al*, 2015; Impens *et al*, 2014; Hendriks *et al*, 2018), likely because the branched peptides that were generated upon protease digest of IRF2BP proteins was too short to be assigned. However, very recently, IRF2BP2 SUMOylation on this conserved lysine residue was found in zebrafish (Wang *et al*, 2019). Taken together, in Hela cells, IRF2BP proteins use a very conserved lysine at their C-terminus for their SUMOylation.

**Figure 2:**
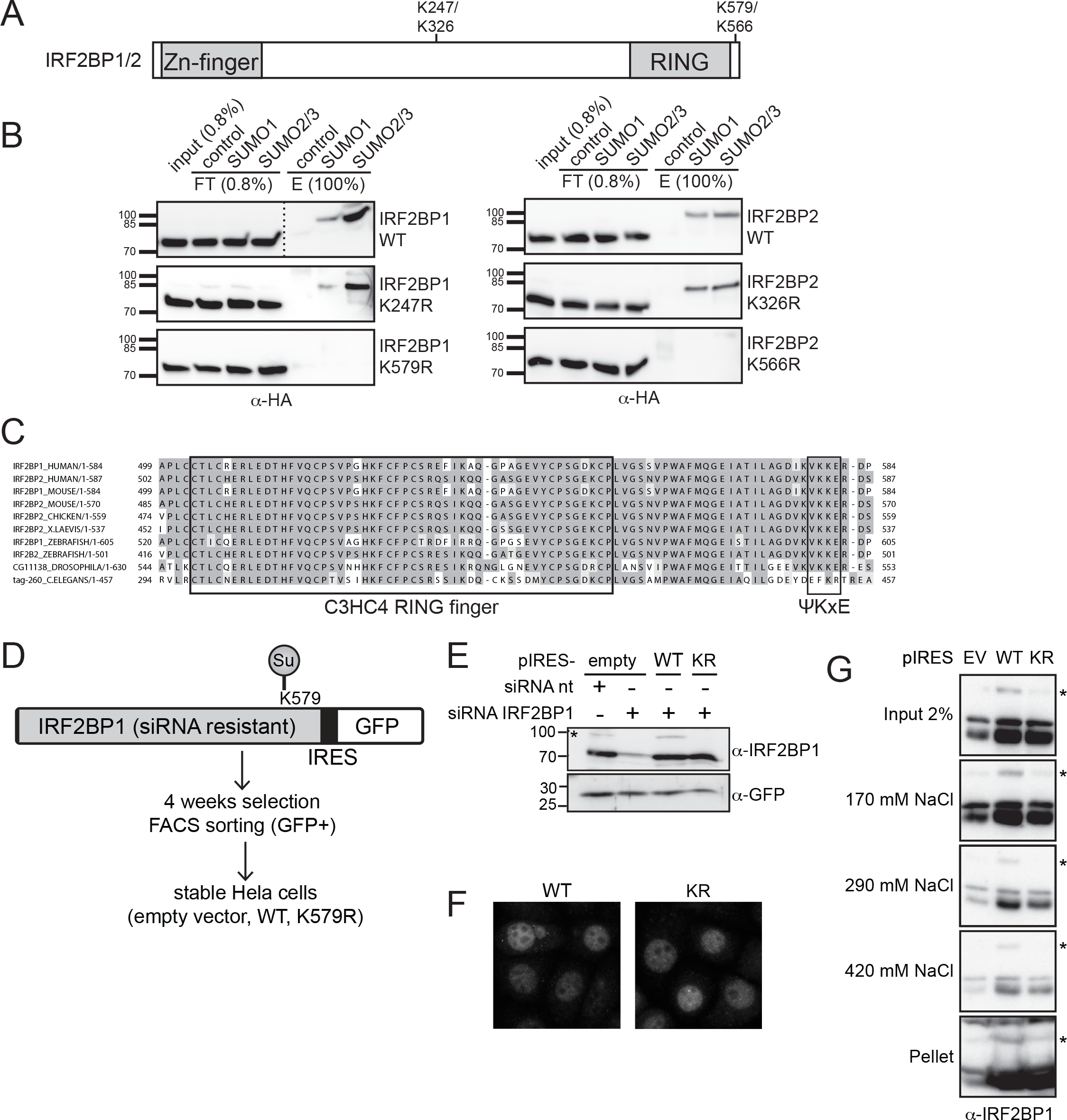
IRF2BP proteins are SUMOylated at their highly conserved C-terminus. (A) Schematic representation of the domain structure of IRF2BP1 and IRF2BP2. The primary sequence suggests two putative SUMO-sites that are conserved among different species. (B) Identification of the endogenous SUMO-sites in IRF2BP1 (left panel) and IRF2BP2 (right panel). Hela cells were transfected with HA-tagged wildtype (WT) or mutant (KR) proteins, endogenous SUMO-IPs were performed and the HA-signal was analyzed by immunoblotting. Mutation of the C-terminal SUMO-site abolished SUMOylation of IRF2BP1 and IRF2BP2 in Hela cells. (C) Clustal omega analysis of IRF2BP1 and IRF2BP2 from *H.sapiens*, *M.musculus*, *G.gallus*, *X.laevis*, *D.rerio*, *D.melanogaster* and *C.elegans* shows high conservation of the C-terminal region including the SUMO-site. (D) Schematic representation of the creation of stable, untagged and siRNA resistant IRF2BP1 WT and K579R Hela cells. Constructs expressing IRF2BP1 variants in an pIRES-hrGFP II (“pIRES”) vector were transfected, selected with antibiotics and FACS sorted for low GFP expression. (E) Stable Hela cells expressing pIRES-empty vector, IRF2BP1 WT or IRF2BP1 K579R were treated with siRNA against endogenous IRF2BP1 or non-targeting (nt) siRNA. Exogenous siRNA-resistant IRF2BP1 was expressed at low levels similar to endogenous IRF2BP1. (F) Wt and mutant IRF2BP1 localizes in the nucleus. After knockdown of endogenous IRF2BP1, stable IRF2BP1 (WT or K579R) cell lines were immunostained for IRF2BP1. Exogenous IRF2BP1 variants show a similar nuclear localization. (G) IRF2BP1 wt and mutant associate with chromatin to a similar extent. Hela cells were lysed in 0.075 % NP40 (Input). After centrifugation, the nuclei were incubated and vortexed with a nuclear extract (NE) buffer containing 170 mM NaCl. The eluates were collected and the procedure was repeated using a NE buffer with higher salt concentrations, first 290 mM, then 420 mM. Wildtype IRF2BP1, the SUMO-deficient K579R mutant and the SUMOylated wildtype form (*) all behave similarly.

### SUMOylation deficient IRF2BP1 cells differ in EGF-dependent transcription

To gain insights into the functional consequences of IRF2BP protein (de)SUMOylation and its role in EGFR signaling, we next generated stable cell lines expressing either IRF2BP1 wildtype or the SUMOylation-deficient mutant. To avoid problems arising from variable expression levels and from tags that were reported to interfere with IRF2BP function (Giraud *et al*, 2014), we generated stable polyclonal Hela cell lines that express untagged and siRNA-resistant variants of IRF2BP1 (wildtype or K579R) using a bi-cistronic vector system (pIRES-hrGFPII) (Fig 2D). GFP selection by FACS was used to specifically select low expressing cells, thus leading to exogenous IRF2BP1 expression that matches endogenous protein levels (Fig 2E).

Expression levels of exogenous wildtype and K579R IRF2BP1 were comparable, suggesting that they have similar stability. Furthermore, localization of exogenous IRF2BP1 was similar in both cell population (Fig 2F), indicating that SUMO does not regulate nucleocytoplasmic transport of these proteins. Another function that has been attributed to SUMO is an influence on transcription factor - chromatin interaction (reviewed, e.g., in (Rosonina *et al*, 2017)).

We therefore compared chromatin binding of wildtype IRF2BP1 with its SUMO-deficient mutant in salt extraction experiments. A significant fraction of IRF2BP1 binds stably to chromatin, and no difference could be observed between wildtype and mutant forms, and also not for SUMOylated wildtype IRF2BP1, which is visible in extracts of stable cell lines (Fig 2G).

To begin to address the question whether SUMOylation of IRF2BP1 may contribute to IRF2BP’s role in gene expression, we next used microarrays to compare the transcriptome between wildtype and mutant cells that were depleted of endogenous IRF2BP1 and grown asynchronously for 48 h in serum containing medium. Indeed, 138 genes were at least 1.5 fold differentially expressed between the two cell lines (see lists of genes in Dataset EV2). Gene Ontology analysis revealed that these genes are involved in the regulation of cell adhesion, proliferation and in the response to growth factor stimuli (Fig EV1).

The most important question was however, whether the lack of IRF2BP1 SUMOylation – and in consequence the lack of temporally controlled deSUMOylation upon EGF stimulation – would result in transcriptional changes. To address this, we again depleted endogenous IRF2BP1 from our polyclonal wildtype IRF2BP1 and K579R cell lines, serum starved the cells for 16 hours and incubated them for an additional hour with or without 100 ng/ml EGF (Fig EV1, Dataset EV2), before cells were harvested and their RNAs quantified using microarray analyses. Consistent with Hela cell studies from Yarden and coworkers (Amit *et al*, 2007), we identified 529 genes significantly regulated by EGF in our stable cell lines (at least 1.5 fold in IRF2BP1 wildtype or in IRF2BP1 KR cells). Intriguingly, for 38 (7%) of those EGF-responsive genes, a significant difference in the amplitude of the transcriptional change induced by EGF could be observed between wildtype and KR cells (at least 1.5 fold, Fig 3A). In light of the early time point (1 hour after EGF treatment), we considered it likely that these 38 genes are directly regulated by IRF2BP1.

**Figure 3:**
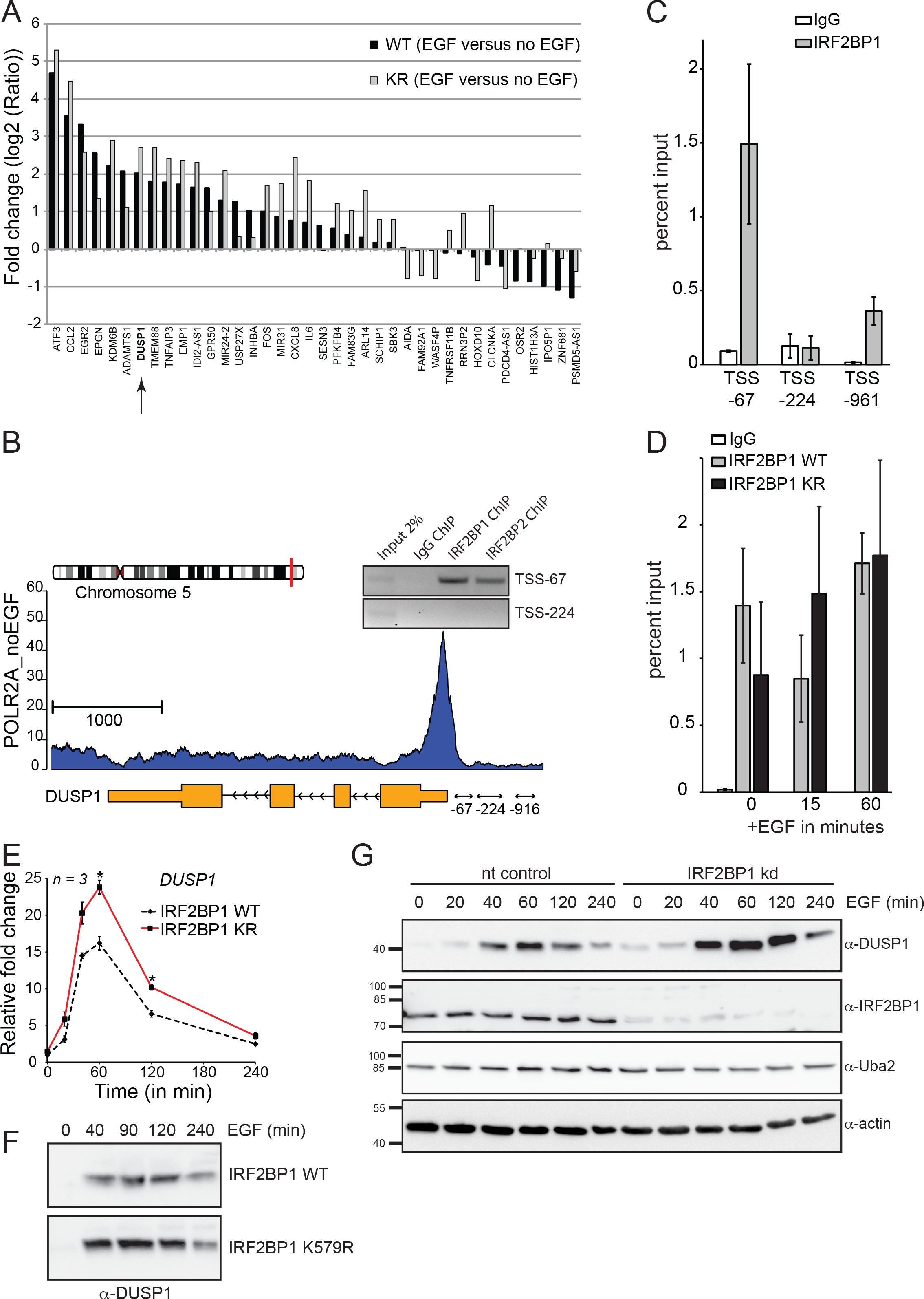
IRF2BP1 deSUMOylation correlates with the transcriptional induction of the immediate early gene DUSP1. (A) Stable IRF2BP1 cell lines (knocked-down for endogenous IRF2BP1) were used to perform a microarray experiment under serum starvation and upon treatment with EGF for 1 hour. A subset of 38 EGF-dependent genes are differentially regulated in IRF2BP1 wildtype compared to IRF2BP1 K579R cell lines (at least 1.5-fold). Among them is the dual specificity phosphatase 1 DUSP1. (B) Chromatin IP reveals association of human IRF2BP1 with the proximal DUSP1 promotor. Gene architecture of human DUSP1. The primers at −243/−67 (“−67”), −473/−224 (“−224”) and −1170/−961 (“−916”) relative to the TSS were used for ChIP experiments. IRF2BP1 and IRF2BP2 bind to the promoter region of DUSP1 between nucleotides −243 and −67. (C) ChIP/qPCR experiments reveal preferential IRF2BP1 binding to the promoter region of DUSP1 between −243 and −67. Data show means +/− SEM from 2 biological replicates. (D) IRF2BP1 wildtype and the K579R mutant both bind the DUSP1 promoter in the absence or presence of EGF. Stable cell lines expressing IRF2BP1 wildtype or K579R were knocked-down for endogenous IRF2BP1, followed by IRF2BP1 ChIP and DUSP1 qPCR of its promoter region −243 and −67. Data show means +/− SEM from 3 biological replicates. (E) Q-PCR data after EGF treatment in the IR2BP1 wildtype and K579R cell lines after knockdown of the endogenous IRF2BP1. IRF2BP1 K579R causes a stronger DUSP1 gene expression compared to wildype. Data show means +/− SEM from 3 biological replicates. (F) Immunoblotting of HeLa lysates at indicated times after EGF treatment reveals enhanced DUSP1 expression in IRF2BP1 K579R cells compared to wildtype. Note that top and bottom panels are from the same blot. (G) Immunoblotting of HeLa lysates at indicated times after EGF treatment reveals enhanced DUSP1 expression upon knockdown of IRF2BP1. Uba2 and β-actin serve as independent loading controls.

### The feedback regulator DUSP1 is a direct target of IRF2BP1

IRF2BP proteins are transcriptional co-regulators that seem to interact with many different transcription factors and co-regulators. In MEL cells, 40% of the IRF2BP2 binding sites were found within 5 kb of the transcriptional start sites (Stadhouders *et al*, 2015). We thus asked whether any of our 38 candidate gene proximal promotors had been identified as binding partners for IRF2BP2 in the mouse study. Indeed, DUSP1 (MKP-1) bound IRF2BP2 in the proximal promotor region (Fig EV2). DUSP1 is a well-known immediate early gene and its gene product, <u>Du</u>al <u>S</u>pecificity <u>P</u>hosphatase 1, is an inhibitor of the MAP kinase branch of EGF receptor signaling and thus an important feedback regulator in EGFR signaling (reviewed in (Liu *et al*, 2007)). In consequence, DUSP1 seemed an excellent candidate to gain mechanistic insights into how IRF2BP1 (de)SUMOylation contributes to gene expression. We thus tested by chromatin IP experiments whether human IRF2BP2 and IRF2BP1 bind to the human DUSP1 promoter in Hela cells. Indeed, both proteins bind to a region (−243 to −67) directly adjacent to the TSS (Fig 3B and 3C), confirming DUSP1 as an excellent model gene for further investigations. SUMOylation can modulate the ability of transcription factors to interact with their target genes (reviewed, e.g., in (Rosonina, 2019)). An obvious question was hence whether transient deSUMOylation of IRF2BP proteins regulates its association with the DUSP1 promotor either positively or negatively. This does not seem to be the case here: first, association with the DUSP1 promotor does not differ significantly between IRF2BP1 wildtype and the K579R mutant, and second, EGF treatment, which causes IRF2BP1 deSUMOylation, does not significantly alter the interaction of wt IRF2BP1 with the DUSP1 promoter (Fig 3D). In conclusion, IRF2BP1 binding directly upstream (within 240 nucleotides) of the DUSP1 TSS is independent of its SUMOylation status and is not altered by EGF.

### IRF2BP1 is a SUMO-dependent repressor of DUSP1

Does transient deSUMOylation of IRF2BP1 contribute to timing and/or the amplitude of DUSP1 mRNA and protein expression after EGF treatment? To address this question, we turned to qPCR experiments (Fig 3E) and to immunoblotting (Fig 3F). Indeed, although DUSP1 mRNA and protein levels showed the expected time course of induction and decline after EGF treatment in wildtype IRF2BP1 and in IRF2BP1 K579R cells, the amplitude of DUSP1 expression was significantly enhanced in the mutant cell line (Fig 3E and 3F). We envisioned two different explanations for this observation: Either, IRF2BP1 is a repressor of DUSP1 gene expression that requires SUMO for its repressor function. Or IRF2BP1 is required for DUSP1 expression, but is inactive as long as it is SUMOylated. To distinguish between these scenarios, we asked whether IRF2BP1 knockdown causes alterations in DUSP1 levels. Indeed, as shown in Fig 3G, knockdown prior to serum starvation and EGF treatment caused a strong increase in the amplitude of DUSP1 induction, indicating that IRF2BP1 is a SUMO-dependent repressor of DUSP1.

Taken together, our findings identify IRF2BP1 as a SUMO-dependent repressor of the immediate-early gene DUSP1 and possibly other EGF-responsive genes. While transient inactivation of IRF2BP1 by deSUMOylation is required for timely expression of DUSP1, both depletion or mutation of IRF2BP1 lead to elevated levels of DUSP1 RNA and protein. Transient inactivation of IRF2BP1 by deSUMOylation thus adds an additional layer of control to the timing and amplitude of EGF receptor signaling.

## Discussion

Following up on the idea that we can exploit comparative quantitative SUMO proteomics to search for novel players in signal transduction, we ideed identified a novel regulatory module in EGF receptor signaling, the simultanous and transient deSUMOylation of five transcriptional co-regulators. These are TRIM24, TRIM33 and the three IRF2BP proteins, the latter of which had not been linked to the EGF receptor pathway. Following up on IRF2BP1 specifically, we could show that it controls EGF-dependent expression of numerous immediate early genes. Using its target gene DUSP1 as proof of principle, we found that IRF2BP1 functions as a SUMO-dependent repressor whose deSUMOylation is required (albeit not sufficient) for EGF-induced DUSP1 expression, and whose reSUMOylation controls the amplitude of DUSP1 expression. With the "SUMO switch" discovered here we add an important element that helps understanding temporal control in immediate-early gene expression.

### How does IRF2BP1 deSUMOylation contribute to DUSP1 expression?

Multiple mechanisms have been described by which SUMO contributes to transcriptional regulation of specific genes ((Hay, 2005; Gill, 2005; Guo *et al*, 2007; Rosonina *et al*, 2017; Rosonina, 2019)). For example, it may enhance the interaction of transcription factors with co-repressors or it may alter the affinity of transcription factors for DNA. Most studies describe SUMO as an inactivating modification or repressive transcription mark, but SUMOylation of gene-specific transcription factors can also be associated with transcriptional activation (Lyst & Stancheva, 2007). The SUMO pathway can thus contribute in manifold ways to the regulation of individual genes, and it is hence not surprising that SUMOylation of transcription factors itself is subject to regulation. An early example is the deSUMOylation of the transcription factor Elk1 upon PMA-induced activation of MAP kinases (Yang *et al*, 2003).

SUMOylation does not seem to alter IRF2BP1’s abundance, localisation, global chomatin association and DUSP1 promoter interaction (Fig 2E-G and Fig 3D). How then does SUMO influence IRF2BP1’s repressive function? Our working hypothesis is that SUMO modulates specific interactions of IRF2BP1 on the DUSP1 promotor. Although identification of the relevant interaction partners is beyond the scope of this manuscript, we can speculate about their nature: DUSP1 is an immediate early gene, which shows strong enrichment of polymerase II at the TSS in serum starved Hela cells (Gardini *et al*, 2014) and whose expression is rapidly induced in response to EGF. The underlying mechanism of transcriptional control is unclear, but could involve regulation of transcription pausing, as has been suggested by (Ryser *et al*, 2004) for DUSP1 in rat pituitary cells or regulation of transcription initiation (Ehrensberger *et al*, 2013). A role for SUMO in transcriptional pausing has recently been suggested in the context of severe heat stress, where globally enhanced SUMOylation serves a protective function (Niskanen & Palvimo, 2017). Publicly available data reveal that the human DUSP1 promoter is bound by numerous transcription factors (ENCODE database, e.g. (Johansson-Haque *et al*, 2008)). Moreover, analysis of rat pituitary cells revealed enrichment of several transcription factors and transcription elongation factors such as NELF and DSIF on the proximal DUSP1 promoter in the resting state (Ryser *et al*, 2004; Fujita *et al*, 2009). Considering that IRF2BP1 sits directly adjacent to the TSS, it may bind to any of these factors or to the RNA Polymerase itself, and it is conceivable that SUMO is required for – or inhibits one of these interactions. This way, SUMO may either contribute to the stability of a paused state, or inhibit transcription initiation. An important question for future studies will thus be the identification of SUMO-regulated interaction partners of IRF2BP1 on the DUSP1 promotor.

### A SUMO brake that prevents erroneous gene activation

It is important to note that IRF2BP1 deSUMOylation in response to EGF is necessary, but not sufficient to drive DUSP1 expression: we do not observe premature expression or expression in the absence of EGF in the IRF2BP1 SUMO-deficient cells. A second EGF-dependent event is required, which may for example be MAP kinase dependent phosphorylation of a transcription or elongation regulator. SUMO thus serves as an important brake that prevents both, erroneous activation and overshooting response (Fig 4, Fig EV3). But how can SUMO restrict DUSP1 expression until it is required, if most IRF2BP1 is not SUMOylated in cells? We can envision two scenarios: either IRF2BP1 is quantitatively SUMOylated on the DUSP1 promoter, but not on many other genes to which it binds. This may for example be accomplished by a promoter-specific binding partner that protects IRF2BP1 from isopeptidases. Alternatively, the whole pool of IRF2BP1 undergoes constant cycles of SUMOylation and deSUMOylation, and EGF shifts the equilibrium to the unmodified form. Irrespective of whether the SUMOylated species contributes to the stability of a paused state or blocks transcription initiation, shortening the lifetime of the SUMOylated species would increase the amplitude of transcription. Either model depends on a signaling event that alters the susceptibility of IRF2BP1 for (de)SUMOylation. Whether this is phosphorylation of IRF2BP1 or one of its binding partners or phosphorylation of a SUMO enzyme is one of many exciting follow-up studies.

**Figure 4:**
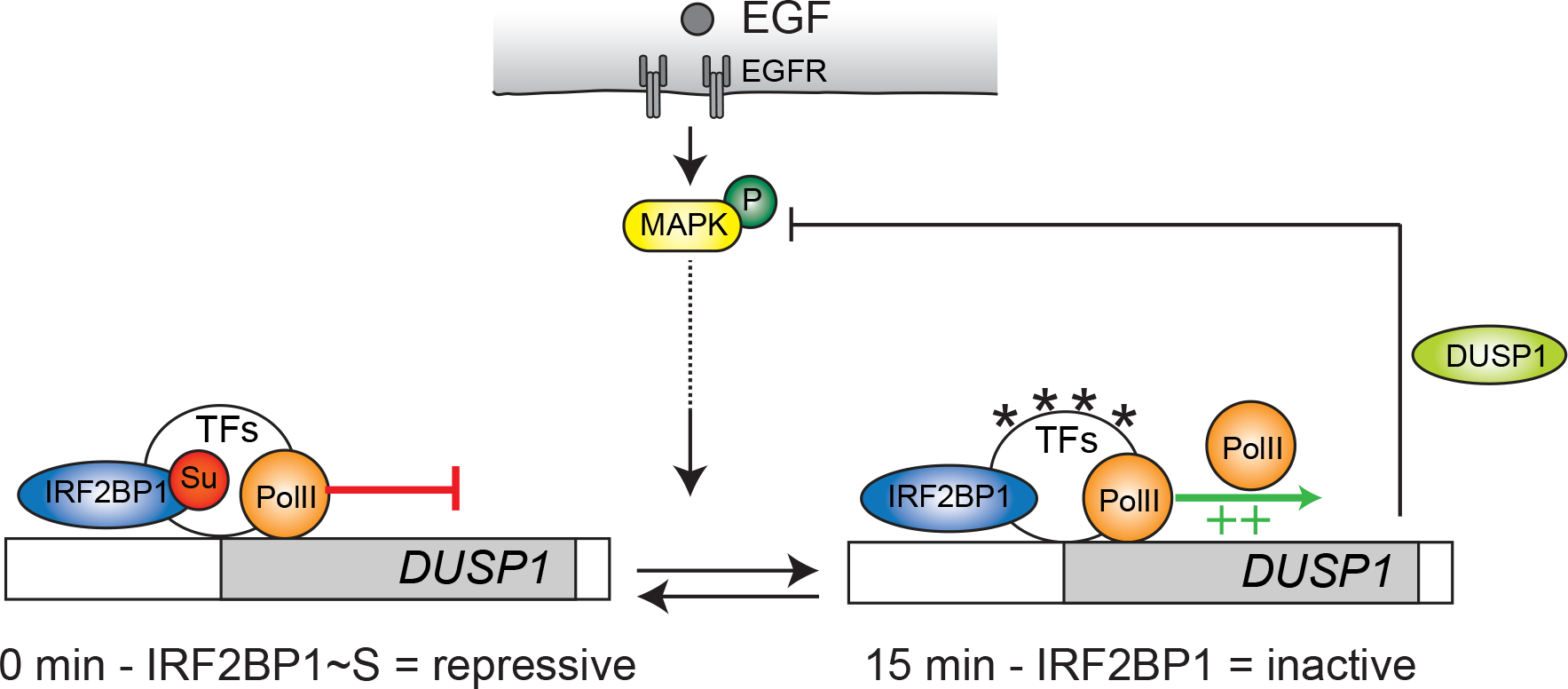
Summary and model. IRF2BP1 is a SUMO-dependent transcriptional repressor of Dual specificity phosphatase 1, a negative feedback regulator of EGFR signaling. Transcription of DUSP1 is repressed by SUMOylated IRF2BP1, which binds to its proximal promotor. EGF receptor signaling yields at least two signals to induce DUSP1 expression, one of which is the transient deSUMOylation of IRF2BP1.

### Outlook

Here we identified an unexpected switch in EGF receptor signaling, the transient deSUMOylation of a small group of transcriptional co-regulators. Following up on one of these, IRF2BP1, we obtained evidence for an important role of this SUMO switch in the temporal control of gene expression, particularly for its direct target DUSP1. It seems very likely that we currently observe only the "tip of the iceberg", as coordinated regulation of the three IRF2BP proteins and the two TRIM proteins is likely to have an even more severe effect on transcription on DUSP1 and other immediate early genes than regulation of IRF2BP1 alone. An important next step will thus be the identification of the upstream event(s) that lead to coordinated deSUMOylation and reSUMOylation of all five co-regulators.

DUSP1 is an important negative feedback inhibitor not just for EGF receptor signaling, but in many pathways that involve MAP kinases (Liu *et al*, 2007). If transient deSUMOylation of IRF2BP and TRIM proteins is an intrinsic signaling event in all pathways that involve MAP kinases, interference with their SUMOylation and/or deSUMOylation may emerge as an interesting strategy to manipulate MAP kinase signaling.

In light of the fact that many transcription factors and regulators are SUMOylated, we speculate that these context-specific "SUMO switches" are frequent elements in signal transduction, particularly in immediate early gene responses.

## Methods

Detailed information on all used resources and methods are provided in the Appendix Supplementary Methods.

### Cells

Hela cells were obtained from Dr. Simona Polo (Milan) and were cultured and treated with 100 ng/ml EGF as described in their corresponding ubiquitin screen (Argenzio *et al*, 2011).

Stable cell lines were generated by transfecting Hela cells with IRF2BP1-pIRES-hrGFPII plasmids (containing IRF2BP1 wildtype or the SUMO-deficient mutant K579R as untagged and siRNA-resistant variants) using polyethyleneimine. After 3 days, cells were selected for one week using G418 and FACS sorted afterwards for low GFP expressing cells (corresponding to low IRF2BP1 expression) on a BD FACSAria Illu™.

### SUMO proteome after EGF treatment

Hela cells were grown for 6-7 doublings in SILAC DMEM medium containing dialyzed FBS, 2 mM L-glutamine and 146 μg/ml lysine and 86 μg/ml arginine. One set of 24 15 cm plates contained “light” lysine and arginine, the other set “heavy” D4-lysine and ^13^C-arginine. Cells were serum starved for 16-18 hours and treated with 100 ng/ml EGF (in PBS-BSA) for 10 min; the other set was treated with PBS-BSA only. For large-scale SUMO-IPs, the cells were lysed in 350 μl 2x lysis buffer per plate and lysates from all 48 plates were combined. SUMO1- and SUMO2/3-IPs were performed as described in (Becker *et al*, 2013; Barysch *et al*, 2014) and samples were analyzed on a LTQ-Orbitrap XL (Thermo Electron, Bremen, Germany). The raw files were then analyzed using Mascot and MaxQuant.

### Microarray analysis

For comparing the RNA expression levels between IRF2BP1 wildtype and the SUMO-deficient KR mutant, Hela cells stably expressing the pIRES-hrGFPII based vectors (IRF2BP1 WT or IRF2BP1 K579R) were treated with siRNA against endogenous IRF2BP1 (GCUUCAAGUACCUCGAAUA[dT][dT] and UAUUCGAGGUACUUGAAGC[dT][dT]) for 72h in a 6-well. After 56 hours, cells were serum-starved for (a) 16 hours, (b) 15 hours and treated for 1 hour with 100 ng/ml EGF, or (c) not serum-starved at all. Efficiency of the siRNA was controlled by using the siRNA transfection mixture on normal Hela cells in parallel, and by checking the protein amounts of IRF2BP1. RNA samples were purified using Nucleospin RNA Plus kit (Machery-Nagel) and gene expression profiling was performed using GeneChip™ HuGene 2.0 ST Array (Affymetrix). Each experiment was done in three independent experiments.

### Chromatin IPs

Hela cells stably expressing pIRES-hrGFPII based vectors (IRF2BP1 WT or IRF2BP1 KR) were treated with siRNA against endogenous IRF2BP1 for 72h on a 10 cm plate. After 56 hours, cells were serum-starved for 16 hours, or for 15 hours and treated for 1 hour with 100 ng/ml EGF. Chromatin IP (ChIP) experiments were performed using the SimpleChIP enzymatic chromatin IP kit (Cell Signalling), exactly following manufacturer’s instructions. The IP was performed with 10 μg of digested chromatin and 2 μg of IRF2BP1 or IRF2BP2 antibodies (Proteintech), or normal rabbit IgG provided in the kit. DNA abundance was quantified by qPCR using the LightCycler® 480 SYBR Green kit and system (Roche).

## Acknowledgements

We thank Dr. Simona Polo for providing Hela cells, antibodies and ideas, and gratefully acknowledge Heidi Ehret, Andrea Frank, Anja Schubert and Ulrike Gern for excellent technical assistance. We thank all members of the Melchior lab for help and useful discussions. This work received funding from the Deutsche Forschungsgemeinschaft (DFG, German Research Foundation) - Project Number 278001972 - TRR 186, the Cluster CellNetworks Postdoc Program (to SVB) and the Peter and Traudl Engelhorn Foundation (to SVB).

## Author contributions

Conceptualization, S.V.B., N.S.V., and F.M.; Methodology, S.V.B., N.S.V., S.K., C.S and H.U.; Investigation, S.V.B., N.S.V., S.K., J.D., T.N.A. and C.S.; Writing – Original Draft, S.V.B. and F.M.; Writing – Review & Editing, S.V.B., N.S.V and F.M.; Funding Acquisition, S.V.B. and F.M.; Supervision, F.M and H.U.

## Data availability

SUMO1 and SUMO2/3 proteome data upon EGF treatment (versus no EGF) are shown in Dataset EV1. The microarray data have been deposited in NCBI’s Gene Expression Omnibus (http://www.ncbi.nlm.nih.gov/geo/) and are accessible through GEO Series accession number GSE135221. Differentially regulated genes in IRF2BP1 WT cell lines versus IRF2BP1 K579R cell lines are also shown in Dataset EV2.

